# Interferon-sensitized hematopoietic progenitors dynamically alter organismal immunity

**DOI:** 10.1101/2024.04.24.590828

**Authors:** Maria Guillamot, Ipsita Subudhi, Varvara Paraskevopoulou, Aleksandr Prystupa, Ikjot Sidhu, Anna Yeaton, Maria Laskou, Carmen Hannemann, Casey Donahoe, Destini Wiseman, Iannis Aifantis, Shruti Naik, Ada Weinstock

## Abstract

Inflammation has enduring impacts on organismal immunity. However, the precise mechanisms by which tissue-restricted inflammation conditions systemic responses are poorly understood. Here, we leveraged a highly compartmentalized model of skin inflammation and identified a surprising type I interferon (IFN)- mediated activation of hematopoietic stem/progenitor cells (HSPCs) that results in profound changes to systemic host responses. Post-inflamed mice were protected from atherosclerosis and had worse outcomes following influenza virus infection. This IFN-mediated HSPC modulation was dependent on IFNAR signaling and could be recapitulated with the administration of recombinant IFNα. Importantly, the transfer of post-inflamed HSPCs was sufficient to transmit the immune suppression phenotype. IFN modulation of HSPCs was rooted both in long-term changes in chromatin accessibility and the emergence of an IFN-responsive functional state from multiple progenitor populations. Collectively, our data reveal the profound and enduring effect of transient inflammation and more specifically type I IFN signaling and set the stage for a more nuanced understanding of HSPC functional modulation by peripheral immune signals.

## INTRODUCTION

Inflammation is a hallmark of infectious, autoimmune, and chronic diseases (Chovatiya and Medzhitov, 2014). Inflammatory reactions are often compartmentalized and driven by localized immune action within tissues. Even in tissue-restricted inflammation, a surge of cellular and molecular mediators with the potential to influence distal responses are released into systemic circulation. While the mechanisms underlying tissue-restricted inflammation have been extensively studied, the long-term consequences of said inflammation, and in particular the enduring impact of focal inflammation on systemic immune function, remains elusive. Such an understanding could elucidate how transient inflammatory encounters, routinely experienced in a discrete tissue location, modify organismal immune function.

Residing in the bone marrow, hematopoietic stem and progenitor cells (HSPCs) hierarchically give rise to all lineages of the blood system (Comazzetto et al., 2021; Wilson and Trumpp, 2006). These long-lived forebears determine organismal immune contexture throughout life. Moreover, both HSPCs and their instructive niche have unique access to factors in the circulation and express an array of receptors to sense circulating cytokine, chemokine, microbial, and metabolic mediators (Pietras, 2017). As such, these cells represent a “central hub” of immunity that is both malleable and exquisitely sensitive to organismal health status. Prior studies have shown that microbial stimuli such as BCG vaccination (Kaufmann et al., 2018; Kleinnijenhuis et al., 2012; Moorlag et al., 2020b), ꞵ-glucan (Mitroulis et al., 2018; Moorlag et al., 2020a; Novakovic et al., 2016), or proinflammatory factors such as oxidized low-density lipoprotein (Bekkering et al., 2014; Groh et al., 2021), interleukin 1 (IL-1) (Moorlag et al., 2018; Pietras et al., 2016), or IL-6 (Tie et al., 2019) sensitize HSPCs. As a result, myeloid-derived progeny of entrained HSPCs exert heightened pro-inflammatory responses, confer protection against secondary infection challenge, and direct persistent alterations to the immune compartment following infection (Cheong et al., 2023; Ciarlo et al., 2020; Kaufmann et al., 2018; Khan et al., 2020). This training effect has been traced to epigenetic rewiring, histone modifications, and the induction of poised chromatin states by transcription factors such as AP-1 and C/EBPb (Cheong et al., 2023; de Laval et al., 2020). HSPCs are thus able to integrate and remember prior inflammatory experiences and confer heightened functionality onto their progeny.

Clinical and experimental data underscore a role for innate immune memory as a key mechanism underlying non-specific and heterologous protection to infectious agents and vaccine responses. Paradoxically, patients with non-communicable inflammatory skin conditions (Narla et al., 2018; Svedbom et al., 2021) and autoimmune disorders (Hsu et al., 2016; Pego-Reigosa et al., 2009) often exhibit innate immune suppression rather than heightened immunity. The sensitization effect of pro-inflammatory stimuli on HSPCs has been widely demonstrated via administration of factors into the peritoneal cavity or systemic circulation. In contrast, tissue inflammation involves both a complex mix and a time-dependent expression of expression of inflammatory signals. The temporal and complex nature of cutaneous inflammatory reactions on HSPCs is less well understood. Recently, the Divangahi group elegantly uncovered that distinct cytokine signals during mycobacterial infection direct HSPC suppression or activation (Khan et al., 2020). This early glimpse of context-specific HSPC function supports a model in which transient signals during active immune responses can have profound and lasting consequences for HSPC-dependent systemic immunity (Cheong et al., 2023; de Laval et al., 2020; Kaufmann et al., 2018; Khan et al., 2020; Mitroulis et al., 2018) and opens the door to understanding the differential impact of signals that are prevalent early in inflammation versus those found at the peak of inflammation.

During local or tissue-restricted inflammation, a myriad of factors are released into systemic circulation and presumably infuse the bone marrow niche (Caiado et al., 2021). Whether and how these factors emanating from sites of discrete inflammation exert long-term influence over HSPCs is unclear. To dissect these facets of localized, complex, and dynamic inflammatory cues that imprint HSPCs and alter subsequent organismal responses, we turned to the skin, a highly compartmentalized organ with a sophisticated immune surveillance system that is routinely subject to focal infections and injuries. The skin is also a prevalent site of inflammatory diseases such as atopic dermatitis and psoriasis that often manifest systemic comorbidities (Korman, 2020). During emergency hematopoiesis, the contribution of bone marrow-derived innate immune cells in infectious and inflammatory reactions remains an area of active investigation (Boettcher and Manz, 2017). We and others have extensively examined long-term imprinting of local skin stem cells by acute inflammation (Larsen et al., 2021; Naik et al., 2017). Conversely, how stem cell compartments in distal organs, and particularly HSPCs, are entrained by skin inflammation remains unaddressed.

Here we employed a well-defined and self-resolving model of imiquimod (IMQ)-induced skin inflammation to examine acute and long-term HSPC responses (van der Fits et al., 2009). We uncovered an unexpected role for type I interferons (IFNs) in rapidly activating HSPCs even prior to overt skin inflammation that subsides long before the peak of skin pathology. Functional studies revealed that post- inflamed mice were more susceptible to influenza virus infection and conversely more protected from atherosclerosis. Bone marrow transplant was sufficient to transfer this immune modulating effect of IMQ- entrained HSPCs. This alteration of organismal immune response could be recapitulated by administering recombinant IFNα intraperitoneally and was lost in mice lacking the receptor for type I IFN (IFNAR) specifically in the hematopoietic compartment, underscoring HSPCs and IFNAR as central mediators of this response. Mechanistically, the immune-modulating effect of type I IFNs could be traced to enhanced chromatin accessibility in LSKs (Lineage^neg^Sca1^+^Kit^+^ progenitors) associated with type I IFN-responsive transcripts and emergent populations of LSKs that were enriched for type I IFN signals. Collectively, these data indicate that the early modulation of the HSPC compartment by type I IFNs in skin inflammation has profound and context-specific effects on systemic immunity. In so doing, we lay the groundwork for future studies to mechanistically delineate how temporal signals early versus late in inflammation, and perhaps even during resolution, alter subsequent organismal responses to infection and inflammation.

## RESULTS AND DISCUSSION

### HSPCs rapidly activate at onset of skin inflammation

To determine how localized skin inflammation impacts the HSPC response, we used a well-characterized model of psoriasis-like skin inflammation (van der Fits et al., 2009). Daily topical application of a TLR7 agonist, IMQ, results in a robust type 17 response that peaks at day 6 of application and, importantly, self-resolves by day 30 (**Fig. 1, A**). Indeed, we observed the onset of skin pathology and the type 17 response starting at day 2 following IMQ application, which peaked at day 6 and entirely resolved by day 30 (**Fig. S1, A and B**).

**Fig. 1.**
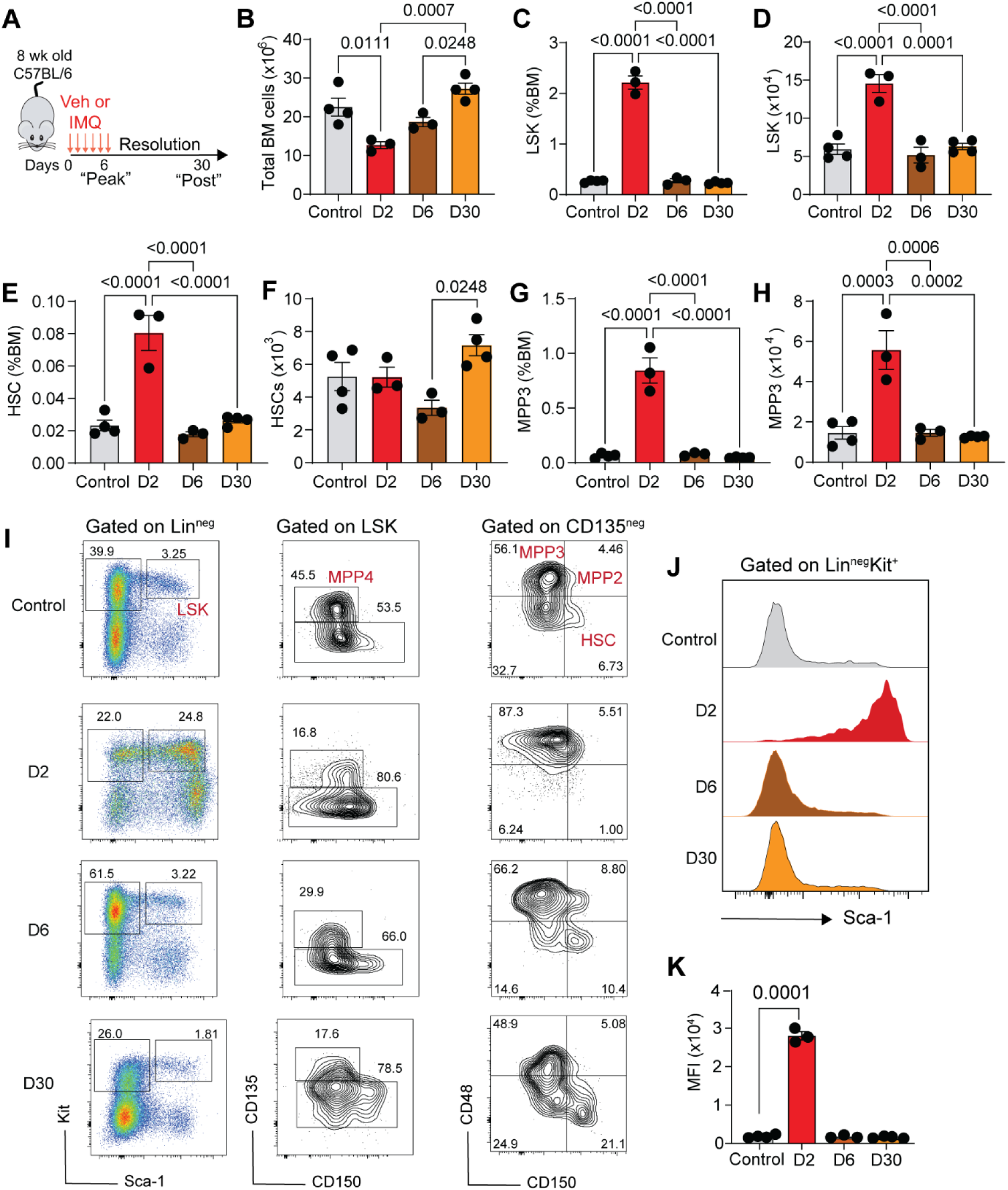
Bone marrow responses rapidly activate and resolve before the peak of skin inflammation. (A) Schematic of experiment analyzing bone marrow (BM) progenitor populations during and after Imiquimod (IMQ) treatment. Analyses were performed 0, 2, 6 and 30 of this protocol (n=3-4 mice/group). (B-H) Absolute numbers and frequencies of bone marrow cells, including (B) all cells, (C-D) Lin-Sca1+c-Kit+ (LSKs), (E-F) Hematopoietic Stem Cell (HSCs) and (G-H) Multipotent progenitors (MPP3). (I) Representative flow cytometry plots and gating strategy. (J) Sca-1 surface expression in LSKs, represented by mean-fluorescent intensity (MFI) and (K) its quantification. P-values were determined via ordinary one-way ANOVA with Tukey’s multiple comparisons test. Plots represent mean ± SEM. Each dot is an individual animal and are representative of two- three independent experiments.

We next assessed the phenotype and response of an array of bone marrow (BM) lineage-negative (Lin^neg^) HSPC, at onset, peak, and resolution of skin inflammation. These include a mix of long-term hematopoietic stem cells (HSCs) that are discerned by LSK markers and are CD135^neg^, CD150^+^, CD48^neg^, and various multipotent progenitors (MPP) that are proximal progeny of HSCs (**Fig. 1, B-I)**. Interestingly, both numbers and frequency of LSKs rapidly increased within 2 days of IMQ treatment and much sooner than the peak cutaneous immune response and pathology in the skin (**Fig. 1, C-D and S1, A- B**). This difference could be traced to a significant increase in the myeloid-biased MPP3s (LSK, CD135^neg^, CD150^neg^, CD48^+^; **Fig. 1, G-H)** at the expense of HSCs (**Fig. 1, E-F**). By day 6, the bone marrow progenitor pool’s composition largely returned to baseline and persisted at these levels throughout the resolution phase at day 30 (**Fig. 1, B-I**). Additionally, the expression of Sca-1, an IFN-responsive activation marker, was highest at day 2 of inflammation and returned to baseline by day 6, indicating that in addition to changes in cellularity, HSPCs were functionally engaged and activated rapidly after inflammation but return to their basal phenotype swiftly (**Fig. 1, J and K**).

Following infection or inflammation, steady state hematopoietic programs are swiftly replaced with a process called emergency hematopoiesis, typified by the rapid generation and swift exodus of innate immune cells from the bone marrow to cope with the looming threat (Caiado et al., 2021). Consistent with the rapid remodeling of the bone marrow progenitor compartment and emergency hematopoiesis, total bone marrow cellularity was reduced at day 2 of IMQ treatment and then returned to baseline by day 6 (**Fig. 1B**). Prior studies administering LPS (Zhang et al., 2016) or β-glucan (Mitroulis et al., 2018) into the peritoneal cavity have found rapid responsiveness of HSPCs, within a day of receiving these systemic microbial triggers. By contrast, here we examined how HSPCs sense localized skin inflammation and found that even in tissue-restricted inflammation, the HSPC compartment is remodeled swiftly. Our data unveil a picture of a highly sensitive progenitor compartment that is rapidly responsive to peripheral inflammation, even before the onset of overt skin disease, and rapidly resolves preceding the peak of skin inflammation.

### HSPCs entrained early in inflammation alter subsequent organismal immune responses

We have previously shown that transient skin inflammation alters local tissue responses through inflammatory training and enrichment of resident type 17 immune cells (Naik et al., 2017). However, if and how transient skin inflammation has subsequent effects on systemic immunity is unclear. Our model of self-resolving skin inflammation allowed us to probe this question using two different (chronic and acute) systemic challenges: influenza virus infection and atherosclerosis. Importantly, these challenges directly impact the lung and cardiovascular system, allowing us to assess systemic immunity without rechallenge in the same tissue. Not only do these disease models plague organs distal to the skin, but each of these models involves a significant innate immune component that allows us to probe functional alterations to the progeny of inflammation-entrained bone marrow progenitors.

We first turned to an acute model of influenza virus infection to assess how responses to a common infectious agent were affected by prior inflammation in a distal location. WT mice were inoculated intranasally with influenza A (strain PR8) after their complete recovery (D30) from skin inflammation (**Fig. 2, A**). Strikingly, while the control group only exhibited a ∼20% mortality rate, 100% of animals in the post-inflamed (PI) group succumbed to the same dose of virus within 10 days (**Fig. 2, B**). PI mice also exhibited significantly more weight loss prior to death (**Fig. 2, C**). Whether increased susceptibility to influenza infection in PI animals is due to improper viral control remains to be seen. Yet, it is tempting to speculate that systemic immune suppression driven by changes to the bone marrow-dwelling progenitors tolerizes innate immune responses in PI mice. Analysis of the bone marrow compartment on day 7 post-influenza infection uncovered significantly fewer LSKs in PI mice when compared to controls (**Fig. S1, F-I**), suggesting an impaired emergency hematopoiesis response may underlie our observed susceptibility to influenza in inflammation sensitized animals.

**Fig. 2.**
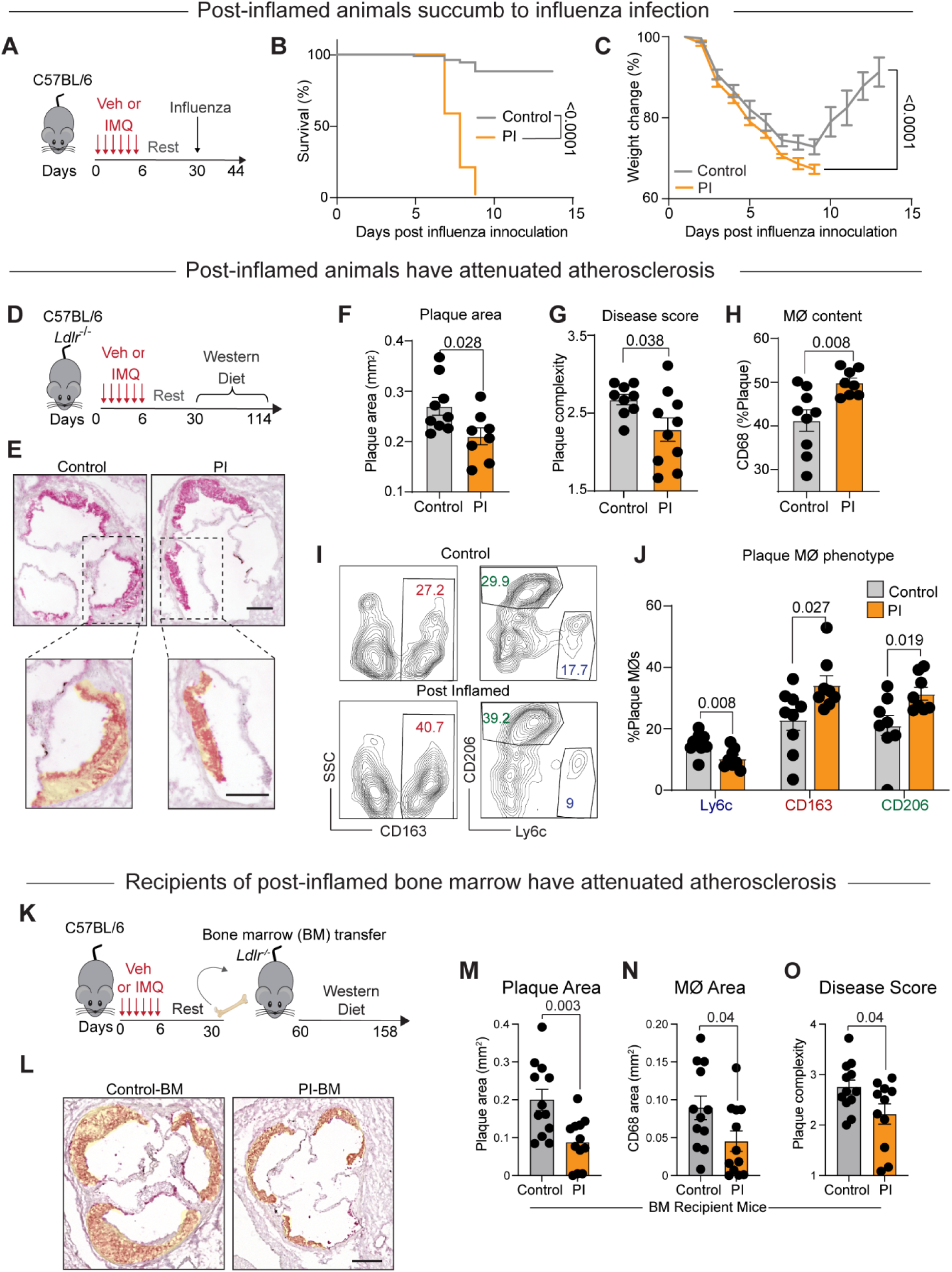
Transient skin inflammation modifies susceptibility to atherosclerosis and influenza infection. (A) Flu infection post-skin inflammation- experimental design (n=13 mice per group). WT mice were treated with IMQ of vehicle for 7 days and rested until full recovery (day 30), after which they were inoculated with influenza and followed for 2 weeks. (B) Survival and (C) weight of mice following flu inoculation at day 0. (D) Atherosclerosis post- skin inflammation- experimental design (n=9-10 mice per group). *Ldlr^-/-^* mice were treated with IMQ or vehicle for 7 days and rested until full recovery (day 30), after which they were fed a Western Diet (WD) for 12 weeks. (E) Representative images of aortic roots stained for CD68. Atherosclerosis plaques are demarcated yellow. Scale bar=0.25mm. (F-H) Morphometric quantification of (F) plaque area, (G) complexity (scored using the Stary scale) and (H) macrophage content. (I-J) Flow cytometry analysis of macrophages from aortic arches, examined for inflammatory (Ly6C) and pro-resolving (CD163 and CD206) markers. Gating strategy presented in (I) and quantification in (J). (K) Atherosclerosis progression in recipients of post-inflamed bone marrow- experimental design. WT mice were treated with IMQ or vehicle for 7 days and rested subsequently until day 30. Bone marrow was then harvested and transplanted to lethally irradiated naive *Ldlr*^-/-^ mice. Four-weeks later, bone marrow recipients started WD feeding for 14 weeks and plaques were assessed thereafter in aortic roots (n=12 mice per group). (L) Representative images (scale bar=0.25mm), (M) plaque and (N) macrophage area, and (O) disease complexity scored using the Stary scale. P-values were determined via unpaired two-tailed Student’s t-test, except for (B) that was determined with Gehan-Breslow- Wilcoxon test. Plots represent mean ± SEM and of two-three independent experiments. Each dot is an individual animal.

To determine if systemic immune modulation by prior skin inflammation was evident in other organs and disease settings, we examined the susceptibility of naïve and PI mice to cardiovascular disease. To induce atherosclerosis, we used mice globally lacking the low-density lipoprotein receptor (*Ldlr^-/-^*), which are prone to developing atherosclerotic plaques when placed on a Western Diet (WD) (Ishibashi et al., 1994). We first confirmed that *Ldlr^-/-^* mice had similar skin pathology and resolution kinetics to WT mice in the primary IMQ response (**Fig. S1, C and D**). *Ldlr^-/-^* mice were treated topically with IMQ or vehicle control for 7 days and rested for 23 days to ensure full recovery of the skin and bone marrow compartment and then placed on a WD to induce atherosclerosis (**Fig. 2, D**). After 12 weeks of WD feeding to allow for plaque development, control and PI animals were assessed for atherosclerosis. In contrast to prior studies indicating heightened immunity following inflammatory training (Bekkering et al., 2014; Cheong et al., 2023; Moorlag et al., 2020a; Moorlag et al., 2020b; Scolaro B, 2023), we found that several disease parameters were lower in PI aortas compared to control. For instance, the plaque’s area and complexity were significantly greater in control versus PI mice (**Fig. 2, E-G**). The aortic plaques in PI mice were less advanced than control mice, as evidenced by Stary plaque complexity scoring (**Fig. 2, G**), and had a larger proportion of macrophages (**Fig. 2, H**), resembling fatty streaks with foam cells and lipid core that arise during early disease development. Macrophages are essential for the development and pathology of atherosclerosis. We therefore characterized macrophage subsets in control and PI aortic arches. PI mice had a lower proportion of Ly6C^+^ inflammatory macrophages and a higher abundance of “pro-resolving” CD163^+^ and CD206^+^ macrophages compared to control plaques (**Fig. 2, I and J**). Importantly, control and PI animals had comparable levels of plasma cholesterol (**Fig. S1, E**), indicating that our identified differences between these groups are not a result of altered cholesterolemia but due to other systemic changes induced by prior skin inflammation.

We next tested if HSPCs alone were sufficient to mediate the immune modulatory effects of transient skin inflammation on organismal responses. To investigate this, we transplanted BMcells from control or PI mice to lethally irradiated, naive *Ldlr^-/-^* recipients. Following transplant, long-lived HSPCs graft into host BM and reconstitute the hematopoietic compartment, providing a system to test the direct contribution of IMQ-entrained HSPCs to systemic immune paralysis. We first confirmed that control and PI donors result in comparable chimerism following irradiation. We then placed hosts on a WD for 14 weeks to induce the atherosclerotic phenotype (**Fig. 2, K**). Remarkably, host mice that received PI BM had significantly smaller atherosclerotic plaques (**Fig. 2, L and M**), with fewer macrophages (**Fig. 2, N**) and overall decreased plaque complexity (**Fig. 2, O**), when compared to control BM recipients. These striking phenotypes were independent of cholesterol levels, which were comparable between the groups (**Fig. S1, J**).

Taken together, these data support the notion that highly localized immune responses in barrier tissues such as the skin can profoundly influence BM progenitors, consequently with enduring effects on systemic immunity. Restraining HSPC function following acute inflammation may represent adaptive organismal mechanisms to prevent exuberant systemic inflammation. On one hand, such immune paralysis is protective in diet-induced atherosclerosis but, on the other hand, renders mice vulnerable to infections. Thus, future studies should explore how long this immune paralysis lasts and the mechanisms that mediate both its endurance and its potential reversion.

### Early and transient type I IFN signaling has lasting impact on systemic responses secondary challenge

IMQ induces a complex inflammatory response, with many proinflammatory cytokines including type I IFNs and IL-17 (Ueyama et al., 2014; van der Fits et al., 2009). In addition, IMQ itself is a TLR7 ligand capable of directly stimulating cells (Hemmi et al., 2002). Mostof these inflammatory factors are enriched at the peak of skin inflammation and not at day 2 when we observed HSPC remodeling (Ueyama et al., 2014). By contrast, type I IFNs are rapidly produced by plasmacytoid dendritic cells within 1-2 days of IMQ application (Wenzel et al., 2005). Indeed, our flow cytometry analysis also revealed a rapid and transient upregulation of Sca-1 on all HSPC cells at day 2 of IMQ treatment (**Fig. 1, J**). Sca-1 (*Ly6a*) is well known to be a type I IFN responsive factor and HSPCs robustly express receptors for these factors (Essers et al., 2009). We thus turned to a publicly available single-cellRNA sequencing dataset of HSPCs to assess the expression of receptors for the aforementioned factors in various BM progenitor populations (GSE108892 (Tikhonova et al., 2019)). This analysis revealed high levels of both type I IFN receptors, IFNAR1 and IFNAR2. on HSPCs, with the highest expression on HSCs and MPP2s (**Fig. 3, A**). There was also a notable expression of an IL-17 receptor (IL-17RA) across various progenitor subsets, but not its key signaling partner IL-17RC, or TLR7, which directly senses IMQ (**Fig. 3, A**). It is possible that IL-17A and TLR7 signaling in the instructive BM niche also contributes to long-term HSPC remodeling and warrants future study. However, given the paucity of IL-17RC and TLR7 on HSPCs and the early Sca-1 responses that corresponded with behavioral changes in HSPCs, we focused our efforts on type I IFNs.

**Fig. 3.**
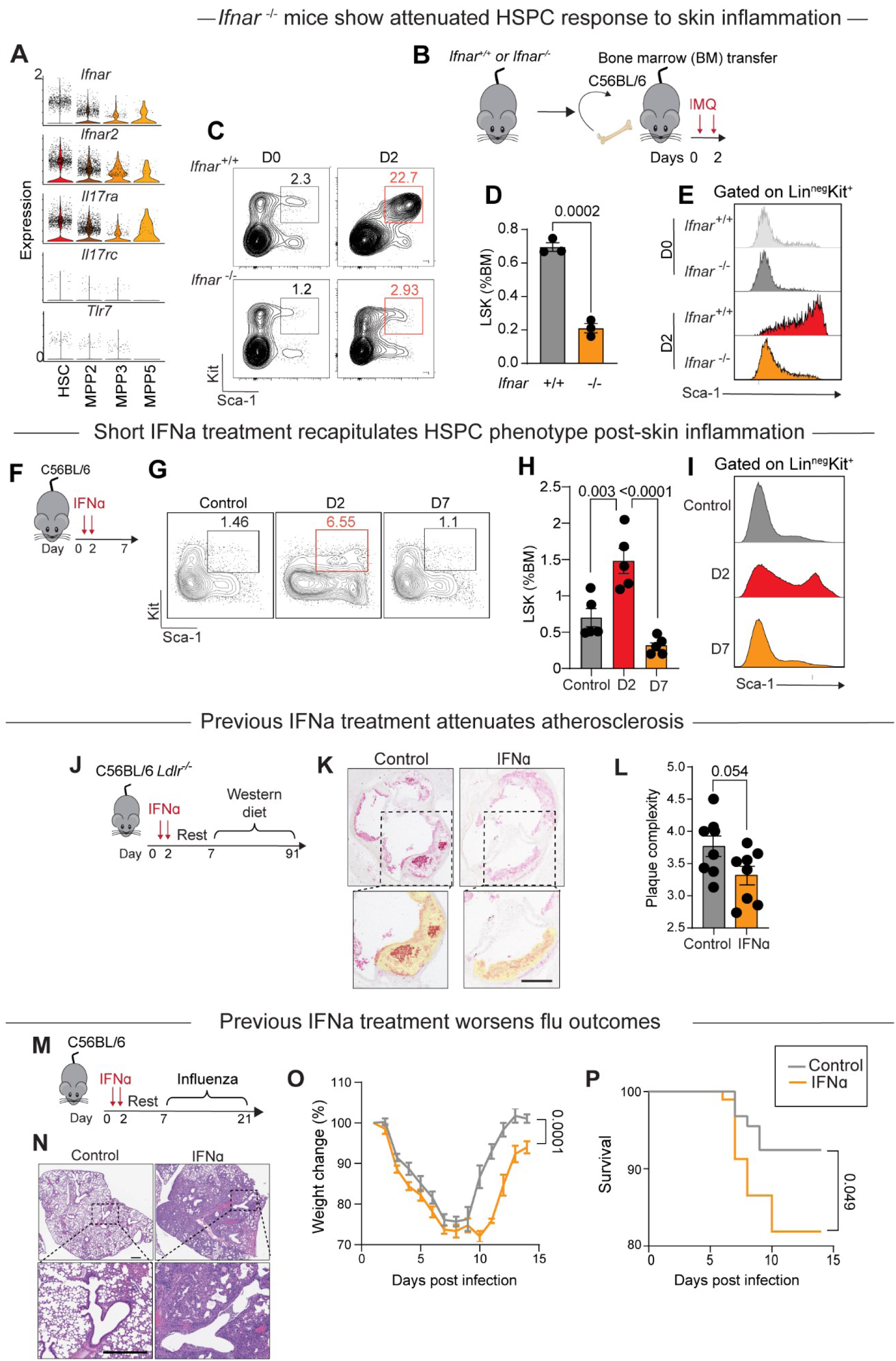
Type I IFN signaling in HSCs drives activation and paralysis with skin inflammation. (A) An available scRNAseq dataset of normal mouse bone marrow (Tikhonova et al.) was analyzed for HSPC expression of cytokine receptors and TLR7. (B-E) *Ifnar*^-/-^ and *Ifnar*^+/+^ bone marrow was transplanted in WT mice. After recovery, recipients were treated with IMQ for 2 days and examined for LSK abundance and Sca-1 expression (n=3/group). (B) Experimental design. (C) Representative flow cytometry plots and (D) their quantification. (E) Sca-1 surface levels in LSKs. (F-I) WT mice were examined for LSK abundance and Sca-1 expression at day 0, 2 and 7 post-injection with IFNα (n=4-5). (F) Experimental design. (G) Representative flow cytometry plots and (H) their quantification. (I) Sca-1 surface levels in LSKs. (J-L) *Ldlr*^-/-^ mice were treated with IFNα for 2 days and rested for 1 week before placing them on WD for 12 weeks (n=8). (J) Experimental design. (K) Representative images of aortic roots stained for CD68. Atherosclerosis plaques are demarcated yellow. Scale bar=0.25mm. (L) Plaque complexity quantified using the Stary scale. (M-P) WT mice were treated with IFNα for 2 days and rested for 1 week before inoculation with Flu. (M) Experimental design. (N) Representative lung images at day 7 post-inoculation. (O) Mouse weights and (P) survival following flu inoculation (n=18). P-values were determined via (D, L, O) unpaired two-tailed Student’s t-test, (H) one-way ANOVA with Tukey’s multiple comparisons test and (P) Gehan-Breslow-Wilcoxon test. Plots represent mean ± SEM and are representative of two-three independent experiments. Each dot is an individual animal.

Prior studies have shown that HSCs express the type 1 IFN receptor (IFNAR) and IFNAR signaling directly activates dormant HSCs, induces strong expression of Sca-1, and exerts long-term effects on myelopoiesis (Essers et al., 2009; Khan et al., 2020). Type I IFNs are robustly produced and systemically secreted early after IMQ inflammation, coinciding with Sca-1 upregulation in LSKs at day 2 of IMQ treatment (**Fig. 1, K-L**) (Griffith et al., 2018). We therefore sought to functionally evaluate the role of type I IFNs in tolerizing HSPCs following skin inflammation. To isolate the IFNAR deficiency to the hematopoietic compartment, we transferred IFN-α/β receptor (*Ifnar*) sufficient and deficient BM to WT hosts (**Fig 3, B**). We treated these BM chimeric mice with IMQ and analyzed the response of immune progenitors. Consistent with our hypothesis that type I IFNs activate HSPCs rapidly in IMQ inflammation, LSKs greatly expanded in WT BM recipients on day 2 following IMQ treatment, whereas *Ifnar^-/-^* BM recipients entirely failed to do so (**Fig 3, C-D**). In addition, progenitors’ Sca-1 expression on day 2 of IMQ treatment was critically dependent on IFNAR signaling (**Fig 3, E**).

We next tested if IFNɑ treatment alone is sufficient to activate HSPCs and the prolonged effects of such transient signaling on systemic immunity. Intraperitoneally injecting WT mice with 50,000U recombinant (r)IFNɑ for 2 consecutive days was sufficient to recapitulate the HSPC activation and resolution kinetics of IMQ treatment (**Fig. 3, F-I**). rIFNα promoted a rapid expansion of the LSK compartment as well as expression of Sca-1 on day 2 (**Fig. 3, F-I**). On day 7 (5 days post-IFNɑ injection), LSK proportions returned to their baseline levels, underscoring the transient nature of IFNɑ-mediated LSK activation. Additionally, this strategy allowed us to assess the specific role of type I IFN modulation of HSPCs and exclude extraneous effects of other cytokines and cell types involved in IMQ-induced inflammation.

Thus, leveraging our transient IFNɑ administration strategy, we tested the causal link between IFNα-mediated HSPC modulation and the long-term systemic immune modulation we had observed post-skin inflammation (**Fig. 2**). Following 2-day intraperitoneal IFNɑ administration, mice were rested for 5 days to ensure resolution of IFNɑ response and then challenged with either atherosclerosis or influenza (**Fig. 3, J and M**). FNɑ primed and control, *Ldlr*-/- mice were fed WD for 12 weeks to induce atherosclerosis (**Fig. 3, J**). Remarkably, the short pretreatment with recombinant IFNɑ restrained atherosclerosis progression, resulting in less complex plaques (**Fig. 3, J-L**). Similarly,7 days after IFNα priming, mice were challenged with influenza infection intranasally (**Fig. 3, M**). Strikingly, mice entrained with intraperitoneal administration of IFNɑ had increased susceptibility to influenza, with severe lung pathology, increased weight loss, and ∼15% lower survival from infection (**Fig. 3, M-P**). This is in contrast to the prophylaxis effects of aerosolized IFNa administered intranasally can be traced to limiting early infectivity in the epithelium, rather than systemic changes to HSPCs (Cheemarla et al., 2021). Similarly, prior upper respiratory tract viral infections can provide heterologous protection by enhancing local anti- viral immunity; this is consistent with our observations that transient priming of HSPCs without engagement of lung immunity results in increased susceptibility to influenza (Cheemarla et al., 2021).

Collectively, these data provide evidence that rIFNɑ was sufficient to phenocopy IMQ-induced alterations to systemic immunity. Here it is important to note that the immune modulatory effect of rIFNɑ was lower than that observed following IMQ-induced skin inflammation. This difference may be due to differences in the amount or duration of IFNɑ administration compared to production following IMQ or due to the contribution of other signals that are present during skin inflammation. Nevertheless, these long-term effects of rIFNɑ alone suggest that rIFNɑ is sufficient to functionally alter HSPCs with enduring consequences for systemic immunity. These findings raise important questions about the difference in transient and chronic type I IFN signaling on HSPC training that warrant further study.

### IMQ imprints HSPCs chromatin and leads to an emergence of a type I IFN progenitor population upon subsequent challenge

Inflammatory imprinting of tissue-generating stem and progenitor cells has been observed in numerous adult stem cell compartments and traced to changes in the epigenetic landscape of these cells (Naik and Fuchs, 2022). In particular, progenitor populations maintain chromatin accessibility at stress response gene loci. This poised chromatin state alters responsiveness to secondary triggers. Given our data thus far indicating that transient programming of HSPC by type I IFNs leads to prolonged effects on systemic immunity, we examined the molecular basis of our observations in two ways: 1) enduring alterations to chromatin accessibility in HSPCs and 2) the alterations to progenitor responsiveness to secondary challenges.

To test the first, we agnostically assessed chromatin accessibility in LSKs purified from control and PI(day 30) mice using Assay for Transposase-Accessible Chromatin sequencing (ATAC-seq; **Fig. 4, A and Fig. S2, A**). We found that a vast majority of accessible chromatin regions were shared between control and PI LSKs (81876, 82.5%) (**Fig. 4, B**) and represent a return to steady state in the PI mice. Moreover, as we are focusing on purified LSKs, these shared chromatin domains also reflect conserved cell identity genes and functions. Importantly, a small fraction of regions was maintained uniquely accessible with either control (5481, 5.5%) or PI (11847, 11.9%) LSKs (**Fig. 4, B)**. A pathway analysis on genes associated with regions of unique accessibility in control identified chromatin-modifying pathways, whereas PI LSKs had unique accessible chromatin in various stress response signaling pathways and type I IFN-responsive genes (**Fig. 4, C)**. These data raise two non-mutually exclusive possibilities for our observed immune alterations: control HSPCs may be more dynamically responsive to systemic challenge, and/or type I IFN-regulated chromatin loci that are enriched in PI LSKs restrain their subsequent function to systemic challenge. A previous study postulated that type I IFN signaling in myeloid progenitors may mediate tolerance to secondary triggers (Khan et al., 2020). These distinct avenues of LSK modulation warrant future investigation and could reveal mechanisms by which transient signals entrain HSPCs.

**Fig. 4.**
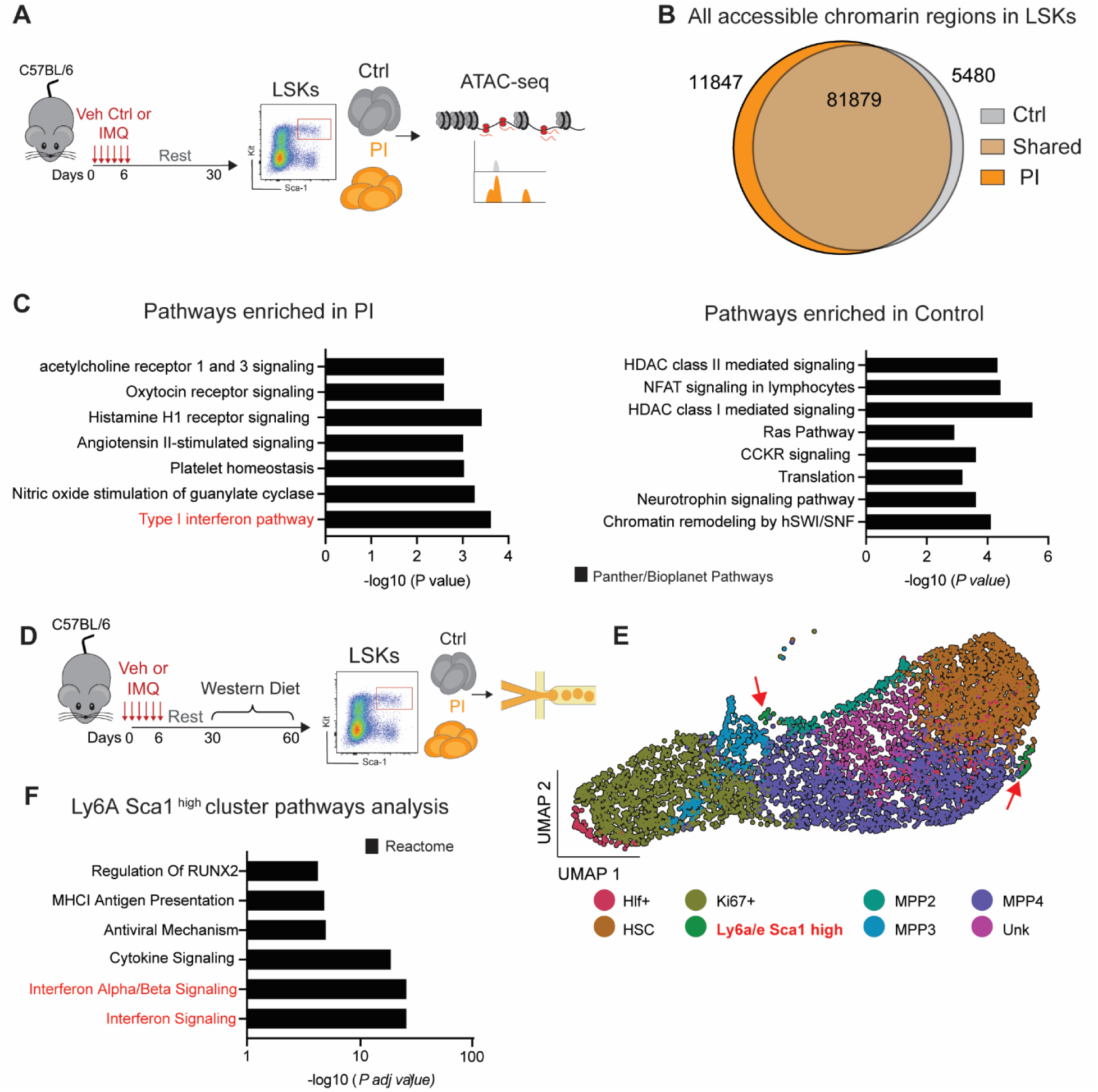
Skin inflammation increases chromatin accessibility in type I IFN genes and promotes a Sca-1^high^ HSPC state. (A) Experimental design. Control and PI (at day 30 post-IMQ treatment and recovery) LSKs were sorted and ATAC-seq performed. (B) ATAC-seq results showing the proportion of accessible chromatin regions that are unique or overlap in the two groups. (C) Pathway enrichment analysis of the unique accessible regions in PI (left) and control (right) LSKs. (D) Experimental design. Control and PI mice (at day 30 post-IMQ treatment and recovery) were fed a WD for 30 days. LSKs were sorted and single-cell RNA-seq performed. (E) Cluster analysis represented in a UMAP. Red arrows indicate Ly6a/e Sca-1^high^ LSKs. (F) Pathway enrichment analysis of Ly6a/e Sca-1^high^ cluster. n=3/group.

We next examined the second facet of immune imprinting, namely altered responses to secondary triggers following inflammatory priming. In this case, we used IMQ as the primary trigger and WD as the secondary challenge. Following 4 weeks of WD, we examined LSK responses from control and PI animals using single-cell RNA sequencing (**Fig. 4, D**). Strikingly, PI mice had a small but significant number of Ly6A+ Sca-1^high^ cells when compared to control LSKs (**Fig. 4, E**). This population was highly enriched for IFN signaling and seemed to emerge from various progenitor subsets, including HSCs, MPP3s, and MPP4s (**Fig. 4, E-F and Fig. S2, B**), suggesting that IFN may simultaneously imprint various progenitor subsets and modify their subsequent function. Future studies should disentangle the relative contribution of IFN imprinting of HSCs versus more committed progenitors (i.e., MPPs).

Taken together, our findings highlight HSPCs as purveyors of the long-term immune paralysis phenotype observed after IMQ-induced training. Though HSPC function in sustaining immunity in health and disease is well appreciated, how these cells are functionally tuned remains less understood. Additionally, whereas a vast majority of innate immune memory research has focused on augmented responses in trained HSPCs, using BM transfers, we find that transient IFN signaling restrains HSPC function, raising the following questions: What is the impact of IFN-mediated progenitor training on differentiated daughter cells? Do differentiated daughter cells exhibit pro-inflammatory cell function or heightened expressions of anti-inflammatory mediators, resulting in compromised immunity to pathogens and immune protection in chronic disease? As inflammatory reactions involve a plethora of mediators, understanding how HSPC responses are modulated in the context of complex milieus and if there is a hierarchy of signals that governs HSPC functionality remains an intriguing question. Understanding the heuristics of inflammatory training of HSPCs, not just in heightening these responses but also in dampening them, as we show here, will open the door to fine-tuning organismal immunity.

## METHODS

### Animals

The following mouse strains were purchased from The Jackson Laboratory: C57BL/6, B6.SJL-Ptprca Pepcb/BoyJ, B6.129S7-Ldlrtm1Her/J. For atherosclerosis studies, *Ldlr*^-/-^ mice were fed WD (100244, Dyets Inc). All animal studies were approved by NYU Grossman School of Medicine’s Institutional Animal Care and Use Committee (IACUC). Mice were bred and maintained at New York University Langone Health Center under specific pathogen-free conditions in Association for Assessment and Accreditation of Laboratory Animal Care (AAALAC)-accredited facilities as well as housed in accordance with procedures outlined in the National Research Council’s *Guide for the Care and Use of Laboratory Animals*. Age- and sex-matched controls were used in all experiments presented, and all experiments were performed in females. Where appropriate, mice were randomly assigned to groups.

### Murine models of inflammation

#### Topical IMQ application

For IMQ studies, seven- to eight-week-old mice in telogen (resting phase of hair cycle) were anesthetized with isoflurane prior to hair removal and IMQ treatment on dorsal skin. 50mg of IMQ cream (5%, Perrigo) was applied to shaved dorsal skin, and application was repeated for up to six days as previously described (van der Fits et al., 2009). Control mice underwent the same procedure of anesthesia and shaving.

#### Atherosclerosis model

Mice were fed WD for the indicated times. Mice were euthanized at the end of each dietary regimen and blood was collected via cardiac puncture, followed by perfusion with saline at physiologic pressure. Hearts were then removed and embedded in OCT (Sakura, 4583; Torrance, CA) and immediately frozen at -80^0^C, while aortic arches were removed and kept in PBS for same day flow cytometry (as described in (Scolaro B, 2023)). Aortic root sections (6 µm) were stained for CD68 (Bio-Rad MCA1957; Hercules, CA) to detect macrophages. Plaque complexity was assessed using the Stary scoring scale (Stary et al., 1995).

#### Influenza infection

Mice were anesthetized with isoflurane and infected intranasally (20 μl) with sublethal 10^5^ EID50 dose (50% embryo infective dose) of the mouse-adapted influenza A H1N1 strain Puerto Rico/34/8 from Charles Rivers (LOT number 4XP201023). Mice that lost more than 30% of their initial body weight were humanely euthanized and considered as non-survivors of this infection.

#### Interferon ɑ treatment

Mice were injected i.p. with 50,000 units of IFNɑ (Miltenyi Biotec, 130-093-130) for 2 consecutive days and rested for an additional 5 days prior to commencing WD or influenza inoculation.

### Bone marrow transplantation

BM cells were harvested from femora and tibias of donor mice. After red blood cell lysis, 2×10^5^ cells were suspended in 0.2 mL PBS and injected retro-orbitally into recipients that were lethally x-irradiated with a total dose of 10Gy. After 4 weeks of recovery and antibiotic treatment, recipient mice were used for subsequent experiments.

### Bone marrow progenitor isolation and quantification

BM cells were harvested from femora and tibias of the indicated mice. After red blood cell lysis, cells were suspended in 0.5 mL PBS+3% FBS and stained for flow cytometry analysis by adding anti-CD16/CD32 antibody (BD Pharmingen; clone 2.4G2) ,anti- CD117 (Biolegend clone 2B8), anti-Sca-1 (Biolegend clone D7), anti-CD34 (BD Biosciences, clones RAM.34), anti-CD135 (Biologend clone A2F10), anti-CD48 (Biolegend, clone HM481), anti-CD150 (Biolegend, clone TC15-12F12), and a BM lineage cocktail for: CD11B (Biolegend clone M1/70), GR-1 (Biolegend clone RB6-8C5), NK1.1 (Pharmingen PK136), Ter119 (Pharmigen), CD4 (Biolegend RM4/5), CD8 (Biolegend) and, B220 (Biolegend clone RA3-6B2). Cells were analyzed using BD LSRFortessa.

### Skin qPCR

Total RNA was extracted from full-thickness skin biopsies RNeasy Plus Micro Kit (Qiagen), and equal amounts of RNA were reverse-transcribed using the superscript VILO cDNA synthesis kit (Invitrogen). Analytes were normalized using the housekeeping gene *Actb*. Primer sequences as follows: *Il17a* (TGACGACCAGAACATCCAGA, AATCGCCTTGATCTCTCCAC), *Il22* ( TGACGACCAGAACATCCAGA, AATCGCCTTGATCTCTCCAC)

### Bioinformatics analysis

Analysis was performed as previously described. Briefly, single-cell RNA sequencing results were analyzed using Seurat (Satija et al., 2015). Samples were demultiplexed with the HTODemux function and negative classified cells were removed. Cells were then further filtered to have between 200-6000 features and less than 10% mitochondrial reads. Samples were processed with the standard Seurat pipeline, including Normalizing and Scaling Data, finding Variable Features, running PCA, Nearest Neighbor, and UMAP. Cells were then clustered at multiple different resolutions, where we identified an optimal resolution of 0.9, identifying 8 distinct clusters. Top markers were found using the presto package (github.com/immunogenics/presto) using the wilcoxauc function, which were then used to annotate cell types. The top 10 most expressed markers from each cell type were used to construct a heatmap. The Seurat FindMarkers function was used to find the top differentially expressed genes between week 4 IMQ-treated samples as compared to week 4 control samples. These genes were then used to do pathway analysis in EnrichR (Kuleshov et al., 2016).

For ATAC-seq analysis, we first removed adaptor sequences and then aligned our data to mm10, mouse reference genome using Bowtie2 package (version - 2.3.4.1 (Langmead and Salzberg, 2012)). Following the removal of duplicate reads, peak calling was performed using MACS2 package (Feng et al., 2012) and the significant peaks output was stored in bed file format. The number of unique and shared accessible regions identified using peak calling in PI and control groups were shown using a Venn diagram (Figure 4B), plotted in an R environment. Differential peak analysis was also performed downstream in an R environment using DiffBind (10.18129/B9.bioc.DiffBind), with PI vs control group comparison and the peaks were filtered to only include significant peaks with FDR cutoff set to (≤0.05) Opening peaks were defined as Fold Change > 0 and Fold Change < 0 for closing peaks. The peaks were further annotated using ChIPpeakAnno (Zhu et al., 2010) package in R where each peak was assigned a gene annotation based on the distance to transcription start site (TSS), with distance cutoff set to 5000 base pairs upstream or downstream of TSS for promotor regions peaks.

The gene annotated peak results were then used for pathway analysis using EnrichR (Kuleshov et al., 2016), and top pathways enriched within Panther and Bioplanet databases were selected for visualization (Figure 4C). Global accessibility heatmaps showing opening and closing peaks in LSK and control samples were generated using deeptools (version 2.0) using computeMatrix function and plotHeatmap with BigWig files and differential peaks results split into opening and closing peaks in PI group as inputs.

### Statistical analysis

GraphPad Prism 9 (GraphPad Software) was used for statistical analysis. Data are expressed as mean ± SEM. Comparison of two groups was analyzed using a 2-tailed student t-test. With three or more groups, statistical analysis was performed using one-way ANOVA, with Tukey’s multiple comparisons testing and Gaussian distribution.

## ACKNOWLEDGEMENTS

We thank Fisher EA, Major J and Weinstock/Naik Lab members for their helpful discussions, advice, and/or critical reading of this manuscript. The following core facilities enabled our study: NYULMC High Performance Computing, Flow Cytometry, Genome Technology Center and Experimental Pathology Core.

## Author contributions

Conceptualization of the study and manuscript/figure preparation (MG, AW, SN), Experimentation (MG, IS, VP, ML, CH, CD, DW and AW), Data Analysis (MG and AW), ATAC-seq and scRNA-seq analysis (AY, IS and AP).

---

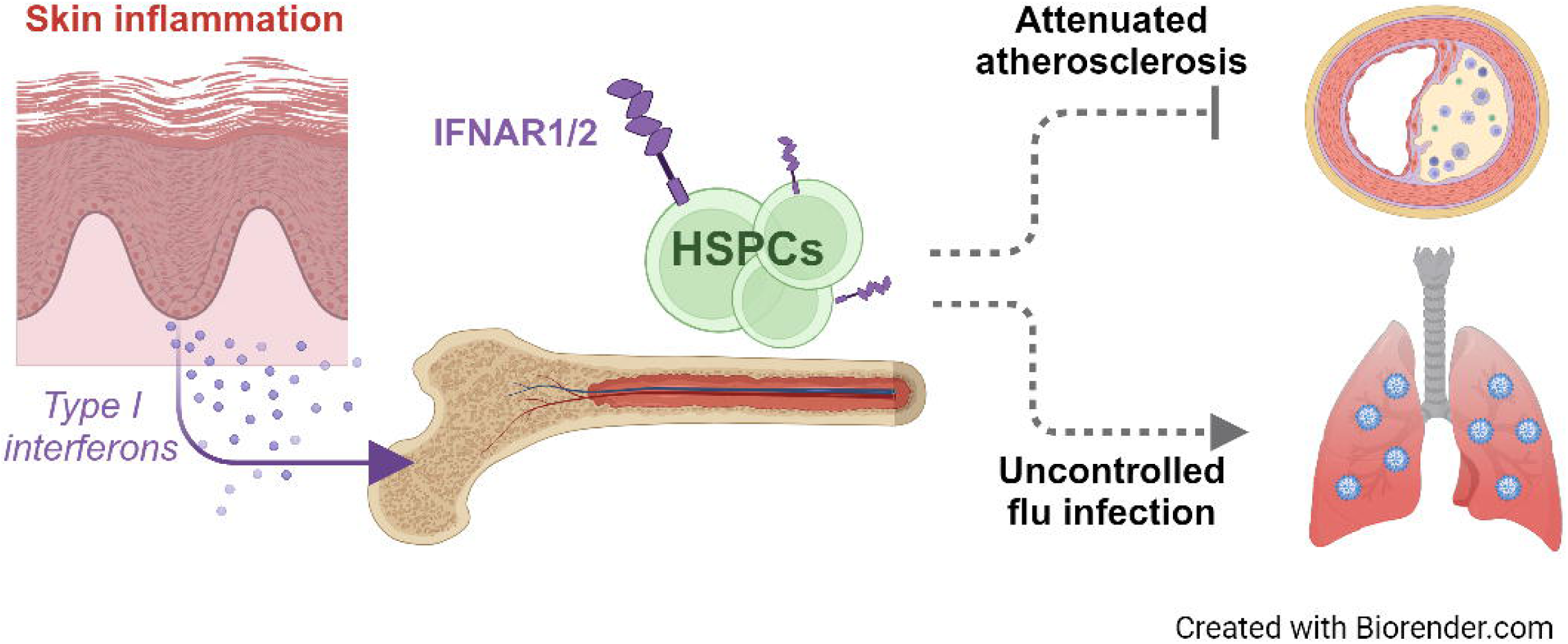

