## Supplementary figures for "Interferon-sensitized hematopoietic progenitors dynamically alter organismal immunity"

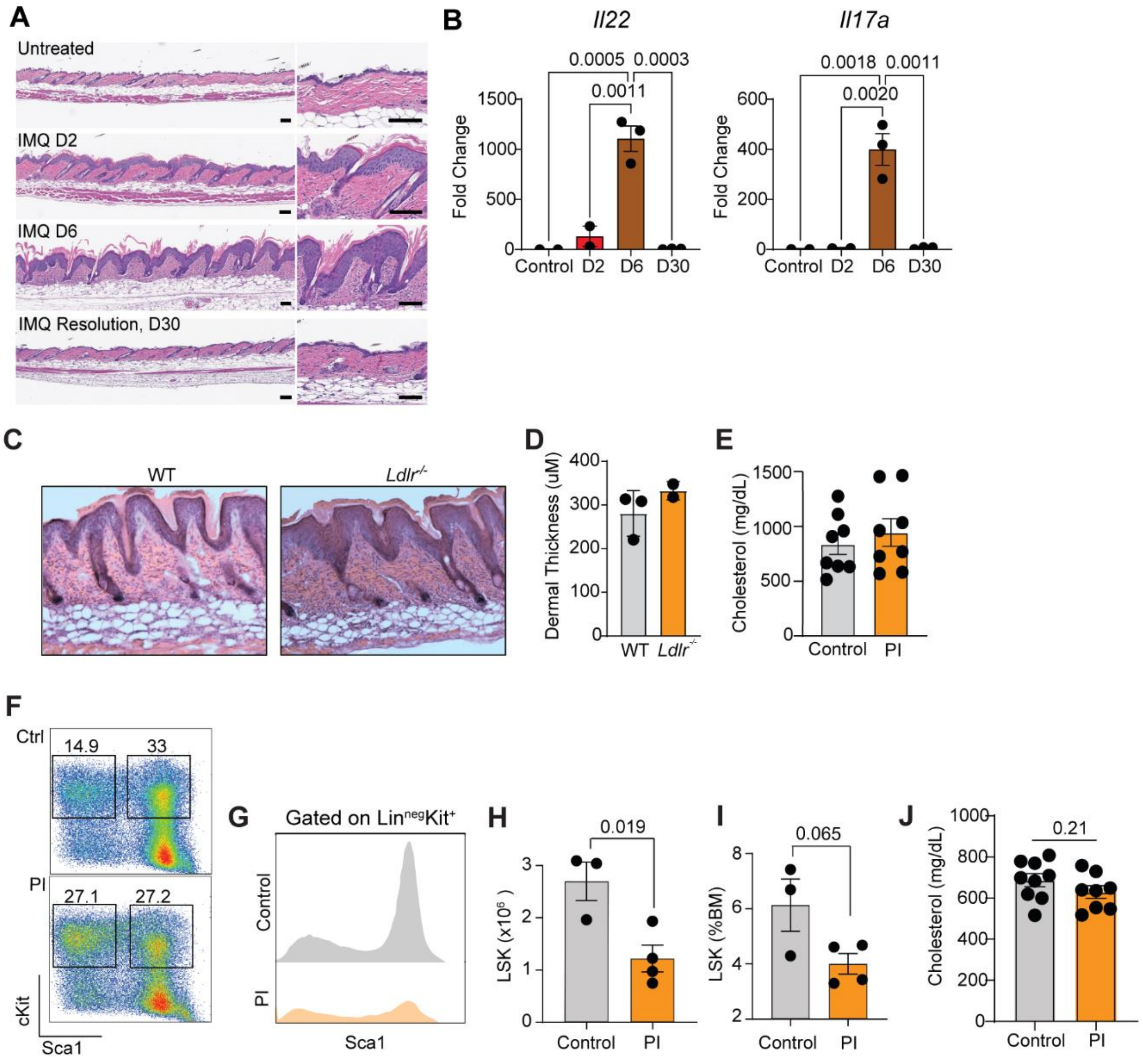


**Fig. S1: Skin inflammation modulates HSPC function**

(A) Skin images following IMQ treatment for 7 days, followed by a recovery period until day 30. (B) Expression of *Il22* and *Il17* in skin at different times following IMQ treatment, relative to untreated mice. (C) Skin images and (D) thickness quantification of WT and *Ldlr*$^{-/-}$ mice. (E) Plasma cholesterol levels in vehicle or IMQ treated *Ldlr*$^{-/-}$ mice after 12 weeks of WD. (F-I) HSPC analysis of control and PI mice at day 7-post influenza inoculation (n=3-4; experimental design presented in Fig. 2H). Flow cytometry representative plots of (F) HSPCs and (G) their Sca-1 levels. (H) Total LSK numbers and (I) their frequency in the bone marrow. (J) Plasma cholesterol levels in bone marrow recipients of vehicle or IMQ treated mice after 12 weeks of WD. P-values were determined via unpaired two-tailed Student’s t-test. Plots represent mean ± SEM. Each dot is an individual animal.

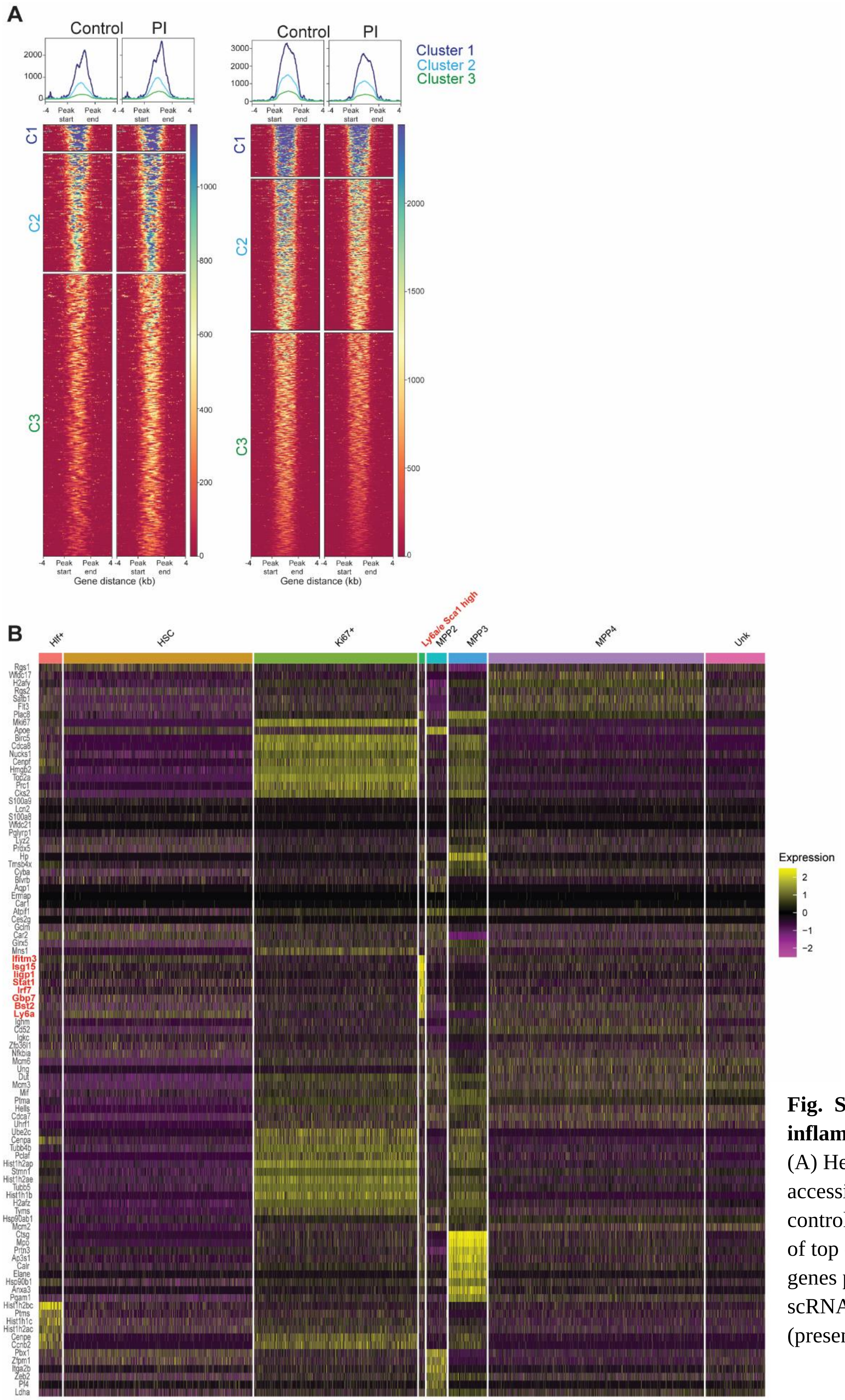


**Fig. S2: Molecular effects of inflammation on HSPCs**
(A) Heatmaps of ATAC-seq accessibility regions enriched in control and PI. (B) Heat maps of top differentially expressed genes per cluster from the scRNAseq experiment (presented in Fig. 4I).
